# Novel Chlamydiae and *Amoebophilus* endosymbionts are prevalent in wild isolates of the model social amoeba *Dictyostelium discoideum*

**DOI:** 10.1101/2020.09.30.320895

**Authors:** Tamara S. Haselkorn, Daniela Jimenez, Usman Bashir, Eleni Sallinger, David C. Queller, Joan E. Strassmann, Susanne DiSalvo

## Abstract

Amoebae interact with bacteria in diverse and multifaceted ways. Amoeba predation can serve as a selective pressure for the development of bacterial virulence traits. Bacteria may also adapt to life inside amoebae, resulting in symbiotic relationships (pathogenic or mutualistic). Indeed, particular lineages of obligate bacterial endosymbionts have been found in different amoebae. Here, we screened an extensive collection of *Dictyostelium discoideum* wild isolates for the presence of such bacterial symbionts using PCR primers that identify these endosymbionts. This is the first report of obligate symbionts in this highly-studied amoeba species. They are surprisingly common, identified in 42% of screened isolates (N=730). Members of the Chlamydiae phylum are particularly prevalent, occurring in 27% of the host strains. They are novel and phylogenetically distinct. We also found *Amoebophilus* symbionts in 8% of screened isolates (N=730). Antibiotic-cured amoebae behave similarly to their endosymbiont-infected counterparts, suggesting that endosymbionts do not significantly impact host fitness, at least in the laboratory. We found several natural isolates were co-infected with multiple endosymbionts, with no obvious fitness effects of co-infection under laboratory conditions. The high prevalence and novelty of amoeba endosymbiont clades in the model organism *D. discoideum* opens the door to future research on the significance and mechanisms of amoeba-symbiont interactions.

## Introduction

As voracious microbial predators, free-living amoebae are important shapers of their microbial communities (Clarholm, 1981). This predatory pressure can alter the presence and abundance of specific microbial constituents in the community (Ronn et al., 2002). Amoeba predation is also postulated to play an important role in microbial evolution, particularly in the evolution of bacterial virulence (Matz and Kjelleberg, 2005; Sun et al., 2018). Bacteria that evolve strategies to avoid or survive amoeba predation would be selected for in amoeba rich environments (Erken et al., 2013). Bacteria that are able to enter amoeba cells (via phagocytosis or other entry mechanisms) but avoid subsequent digestion gain access to an attractive intracellular niche. A diverse collection of intracellular bacterial symbionts of amoebae has been found, some of which appear pathogenic, neutral, or beneficial for their amoeba host (DiSalvo et al., 2015; Jeon, 1992; König et al., 2019; Maita et al., 2018; Shu et al., 2018; Taylor et al., 2012). Some of these symbionts are important human pathogenic species such as *Mycobacteria, Legionella,* and *Chlamydia* (Boamah et al., 2017; Cardenal-Muñoz et al., 2018; Drancourt, 2014; Gomez-Valero and Buchrieser, 2019; Paquet and Charette, 2016; Tosetti et al., 2014).

Bacterial endosymbionts of amoebae were first discovered in the diverse, free-living amoeba genus *Acanthamoeba* (Horn and Wagner, 2004). These amoebae are found in soil and freshwater environments and can protect themselves and their bacterial associates by forming cysts under environmentally unfavorable conditions (Rodríguez-Zaragoza, 1994). The presence of such endosymbionts was known long before they could be specifically identified because the inability to isolate obligate associates from their hosts made characterization challenging. Once molecular methods became available, it was clear that these bacteria were from specific lineages in the Chlamydiae, *Proteobacteria*, *Bacteriodetes,* and *Dependentiae* (Delafont et al., 2015; Horn and Wagner, 2004; König et al., 2019; Maita et al., 2018; Samba-Louaka et al., 2019; Schmitz-Esser et al., 2008). These endosymbionts interact with their amoeba hosts in varied ways. Some inhabit host-derived vacuoles, while others persist in the cytoplasm. A *Neochlamydia* symbiont can protect its host against the detrimental consequences of *Legionella* bacterial infection (Ishida et al., 2014; König et al., 2019; Maita et al., 2018) and other strains have been shown to improve the growth rate and motility of their hosts (Okude et al., 2012). Other known amoeba endosymbionts are from the less-studied genera *Amoebophilus* and *Procabacter*. Genome sequencing of the amoeba symbiont *Candidatus Amoebophilus asiaticus* found no evidence that it could provide novel pathways to supplement host nutrition, but other fitness affects have not been tested (Schmitz-Esser et al., 2010).

Overall, a high proportion of sampled *Acanthamoeba* (25-80%) are infected with endosymbionts and these serve as useful models for understanding how amoebae can serve as training grounds for bacterial intracellular adaptation. The broader distribution of symbiotic bacteria among other amoeba species and their prevalence in natural host populations, however, is unclear (Schmitz-Esser et al., 2008). The potential of amoebae to harbor such bacteria could be of concern to human health. For instance, environmental Chlamydiae have been associated with respiratory diseases and adverse pregnancy outcomes in humans and cattle, although the extent of their importance in causing animal illness is unknown (Borel et al., 2018; Greub, 2009; Taylor-Brown et al., 2015; Taylor-Brown and Polkinghorne, 2017).

Another amoeba that has been used as a model to study host-bacteria interactions is the social amoeba, *Dictyostelium discoideum*. This amoeba, distantly related to *Acanthamoeba*, is terrestrial, living and feeding as single cells in forest soils while the bacterial food supply is plentiful. When food becomes scarce, the amoebae begin a social cycle, which entails thousands of cells aggregating and forming a slug to crawl to the surface of the soil. The cycle culminates in the formation of a fruiting body, made of a ball of spores (sorus) sitting on top of a stalk, which can then be dispersed to more favorable conditions (Kessin, 2001). In many cases, bacteria are cleared from *D. discoideum* cells during the social cycle through innate immune-like mechanisms (Chen et al., 2007). However, vegetative amoebae can be infected with a range of human pathogens such as *Mycobacterium, Bordetella*, and *Legionella* (Bozzaro and Eichinger, 2011; Taylor-Mulneix et al., 2017). As a model system *D. discoideum* has been fruitfully utilized to describe the role of host and pathogen factors in mediating infection mechanisms and virulence (Annesley and Fisher, 2009; Bozzaro et al., 2019; Bozzaro and Eichinger, 2011; Cosson and Soldati, 2008; Steinert, 2011; Steinert and Heuner, 2005).

Recently, a variety of bacterial symbionts have been discovered to colonize *D. discoideum* (Brock et al., 2018; DiSalvo et al., 2015; Haselkorn et al., 2018). The *D. discoideum* microbiome is composed of both persistent and transient bacterial associates (Brock et al., 2018). Three different newly named *Paraburkholderia* species, *P. agricolaris*, *P. hayleyella*, and *P. bonniea* are persistent bacterial symbionts that intracellularly infect vegetative amoebae and spore cells (Brock et al., 2020, 2011; Haselkorn et al., 2018; Khojandi et al., 2019; Shu et al., 2018). While these *Paraburkholderia* themselves do not nourish *D. discoideum*, they confer farming to their hosts by allowing food bacteria to survive in the fruiting body. These food bacteria can then be re-seeded in new environments during spore dispersal, which benefits hosts in food-scarce conditions (Brock et al., 2011; Haselkorn et al., 2018; Khojandi et al., 2019). In natural populations, an average of 25% of *D. discoideum* are infected with *Paraburkholderia* (Haselkorn et al., 2018). Other bacteria, some edible and others not, have the ability to transiently associate with *D. discoideum* even in the absence of *Paraburkholderia* co-infections (Brock et al., 2018). Thus, natural *D. discoideum* hosts a small-scale microbiome and is gaining traction as a simple model system to study the mechanisms of microbiome formation (Dinh et al., 2018; Farinholt et al., 2019). However, all previous sampling of natural *D. discoideum* associates has relied on isolating bacterial colonies from nutrient media plates, which would miss any bacterial species, such as obligate amoeba symbionts, incapable of growing under these conditions.

Since obligate endosymbionts are common in other free-living amoebae, we endeavored to identify them in *D. discoideum.* Here, we screened a frozen stock collection of 730 natural *D. discoideum* isolates. We first used a 16S rRNA gene amplification strategy with “universal” primers and direct sequencing to detect and identify bacteria within *D. discoideum* spores in a single population. In addition to detecting expected *Paraburkholderia* symbionts, we identified many Chlamydiae and *Amoebophilus* sequences, and a few *Procabacter* sequences in screened samples. To determine the extent of Chlamydiae and *Amoebophilus* symbionts in our collection we then used symbiont specific PCR screening. From each of these prevalent bacterial lineages, we found genetically distinct novel species and were able to visualize representative endosymbiont cells within spores via electron microscopy. We also used antibiotics to cure natural hosts of Chlamydiae and *Amoebophilus* and compared host fitness to uncured counterparts. Finally, we looked at the fitness effects of co-infections since co-infections occur in nature.

## Results

### Obligate endosymbionts are common in wild-collected clones of *Dictyostelium discoideum*

To investigate the presence and prevalence of obligate bacterial symbionts in *D. discoideum*, we screened a frozen collection of 730 natural *D. discoideum* isolates for bacterial DNA. The vast majority of our screened isolates were collected from the Eastern half of the United States (Fig. 1a) and had previously been screened for *Paraburkholderia* (Haselkorn, *et al*. 2018, Table S1). This included 16 populations with seven or more individuals collected. Our largest collection from Virginia consists of about 200 individuals collected in 2000 and in 2014. Across all locations we found over 41% of samples harbored at least one obligate endosymbiont. Specifically, 117 (∼16%) were infected with *Amoebophilus* and 198 (∼27%) with Chlamydiae. Sixteen (2.1%) were co-infected with both *Amoebophilus* and Chlamydiae (Fig. 1b). Two isolates (both from Virginia) were infected with an obligate bacterium identified as belonging to the genus *Procabacter*.

**Figure 1.**
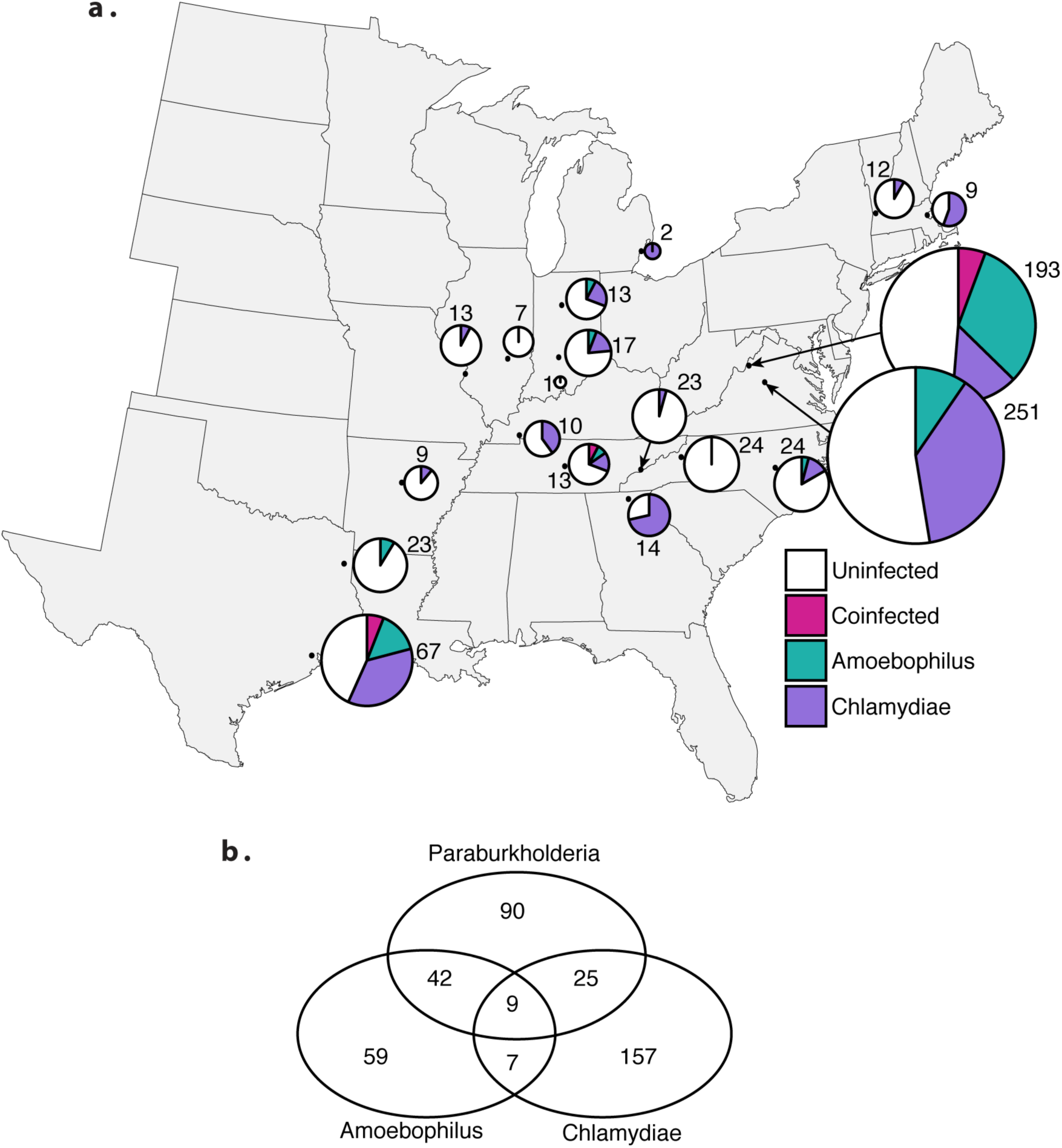
Unculturable symbiont prevalence in natural *D. discoideum* populations. (a) Pie charts indicate Chlamydiae, *Amoebophilus*, and Chlamydiae*-Amoebophilus* co-infection patterns in respective populations of *D. discoideum* isolates from across the Eastern United States. *Procabacter*, due to its low prevalence, is not included. Numbers next to pie charts indicate the number of isolates for each population. (b) Of all symbiont positive amoebae isolates, several are co-infected with more than one symbiont species, represented by the Venn diagram indicating the number of isolates infected with *Paraburkholderia, Amoebophilus,* and Chlamydiae.

### The *Procabacter, Amoebophilus*, and Chlamydiae endosymbionts of *D. discoideum* are novel and phylogenetically distinct from known symbionts

For *Procabacter*, *Amoebophilus*, and the three most prevalent Chamydiae strains (those occurring in more than one individual) we sequenced the full length 16S rRNA for our *D. discoideum* endosymbionts to better determine their evolutionary relationships. In all cases, the 16S rRNA haplotypes were novel and highly diverged, ranging from 6-12% sequence difference from other sequences currently in GenBank or SILVA (Table S2). For the *Procabacter* and *Amoebophilus* endosymbionts, only a single haplotype was found for each. The *Procabacter* endosymbiont of *D. discoideum* is sister to all of the *Procabacter* endosymbionts of *Acanthamoeba* (Fig. 2a). While it is most closely related to this genus, forming a clade with 100% bootstrap support, the 16S rRNA sequence is only a 93% match. A similar pattern is seen for the *Amoebophilus* symbiont of *D. discoideum* (Fig. 2b.). This symbiont groups most closely with the putative *Amoebophilus* symbiont from genome assembly of the recently sequenced *Dictyostelium polycephalum.* This Dictyostelid symbiont clade is sister to the *Amoebophilus asiaticus* endosymbionts of *Acanthamoeba*, and is 6% diverged at the 16S rRNA gene (Table S2).

**Figure 2.**
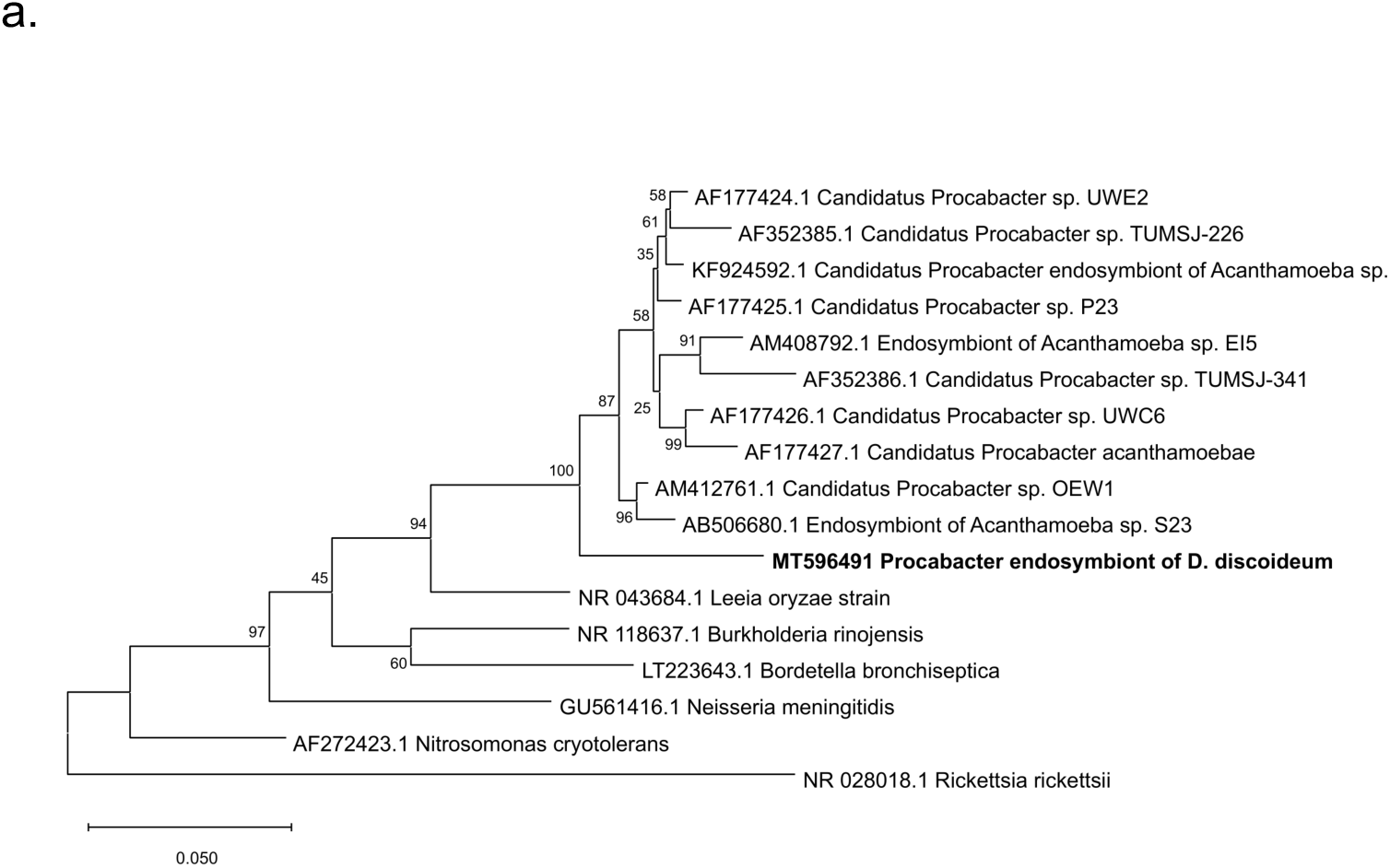
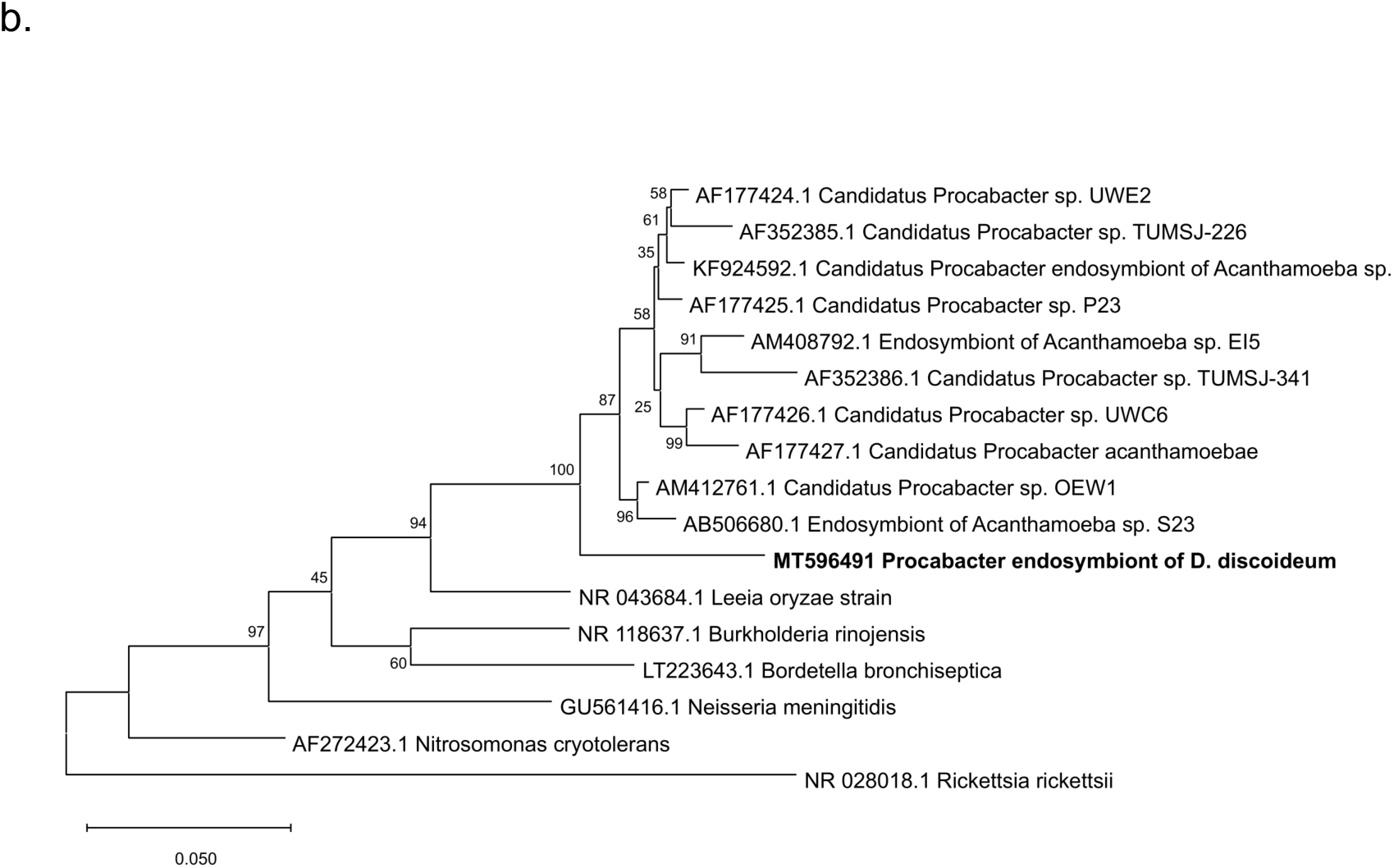
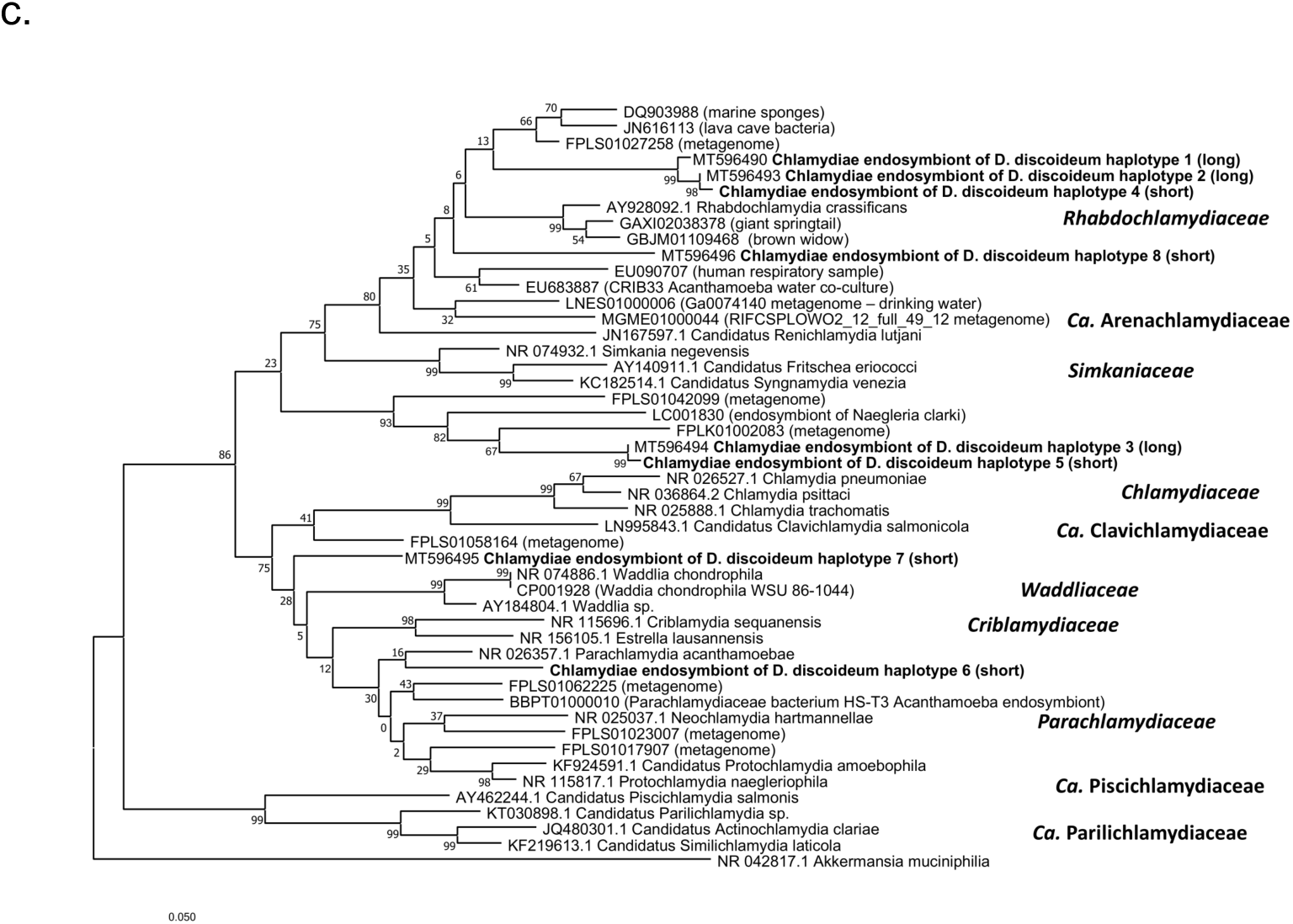
Unculturable endosymbionts of *Dictyostelium* are related to unculturable endosymbionts of other amoebae. 16S rRNA phylogenies of *Procobacter* (a) *Amoebophilus* (b) and Chlamydiae (c & d) endosymbionts constructed using the Maximum Likelihood method in Mega7, with 1000 bootstrap replicates. Family names, where applicable are delineated in bold italics.

There are 8 distinct and novel haplotypes of *D. discoideum* endosymbionts in the Chlamydiae. For the three most prevalent haplotypes (1, 2, and 3), we sequenced the full 16S rRNA gene to reconstruct a more resolved phylogeny which included their nearest neighbors from the SILVA database and representatives from many Chlamydiae families (Fig. 2c). There are currently 6 formally named and 8 proposed Chlamydiae families (Pillonel et al., 2018; Taylor-Brown et al., 2015), and several other recently discovered novel clades from marine sediments (Dharamshi et al., 2020). The *D. discoideum* Chlamydiae endosymbionts do not appear to be closely related (<97% sequence match) to any of these groups. Haplotypes 1 and 2 and 4 form their own well-supported clade, grouping with the *Rhabdochlamydiaceae* and *Ca.* Arenachlamydiaceae, sister to the *Simkaniaceae*. Many of the internal nodes on this phylogeny have low bootstrap support, making more detailed phylogenetic inferences impossible. This *D. discoideum* endosymbiont clade, however, does have only a 91% identity to its closest phylogenetic neighbor and may represent a new genus or family of Chamydiae. Greater than 10% sequence divergence at the full length 16S rRNA gene has been proposed as family level difference (Everett et al., 1999), although full genome sequences may be necessary to make this assertion. Detailed searches for this most common Chlamydiae haplotype 1 in the IMNGS database returned 13 matches (>99% sequence identity), mostly as “uncultured bacteria” from soil metagenomic studies (Table S3). One such study was in the Northeast United States (Massachusetts), in a region close to where we sampled *D. discoideum*, and it is likely that amoeba endosymbionts are being sequenced as part of these metagenomes.

Our other Chlamydiae haplotypes are scattered throughout the phylogeny. Haplotypes 3 and 5 form a well-supported clade and may be sister to *Rhabdochlamydiaceae*/ *Ca.* Arenachlamydiaceae/ *Simkaniaceae* families. These haplotypes have a greater than 11% nucleotide difference from its nearest neighbors on the phylogeny (and in the SILVA and NCBI databases, Table S2) and may represent a novel family as well. Haplotypes 6, 7 and 8 fall in different places in the phylogeny and do not group closely with any other taxa, although these haplotypes are based on only 220bp of sequence information so their taxonomic identity is less clear. Finally, while the newly discovered marine Chlamydiae lineages (Dharamshi et al., 2020) were not included in this phylogeny, when we aligned our Chlamydiae haplotypes with these sequences, the closest match had only an 89% sequence identity.

### Distribution of endosymbionts across populations

While the average overall prevalence of infection of Chlamydiae endosymbionts was 27%, it varied among populations, ranging from 0% to 75% (Table S1). The *D. discoideum* Chlamydiae haplotypes 1, 2 and 3 were the most prevalent in natural populations. Haplotype 1 represented 79.8% of all *D. discoideum* Chlamydiae infections, while haplotype 2 was 13.6% and haplotype 3 was 3.5%. Haplotypes 4-8 were represented only once. Haplotype 1 was found in 12 populations, haplotype 2 was found in 8 different populations, and haplotype 3 was found in three populations. The diversity of Chlamydiae strains was the highest in the Virginia Mountain Lake collection from 2000, which had six of the eight haplotypes (1-6), and the second highest diversity of Chlamydiae haplotypes was found in the Texas Houston Arboretum collection (1, 2, 3 and 8). Interestingly, haplotype 1 was the only haplotype circulating in the most recent 2014 Virginia Mountain Lake collection. While haplotype diversity was lower in this collection, Chlamydiae prevalence (for all haplotypes) in the region increased from 20% in 2000 to 38% in 2014.

The one strain of *Amoebophilus* found to infect *D. discoideum* was distributed across 9 populations, although at low prevalence, often with just a single individual infected in many locations. This endosymbiont was generally less prevalent than Chlamydiae, with the exception of the 2000 Virginia Mountain Lake population, where it infected 37% of the population. *Amoebophilus* prevalence in this region, however, decreased to 10% in 2014. *Amoebophilus* was also relatively prevalent in the geographically-distant Texas Houston Arboretum population, infecting 23% of the population.

### Obligate endosymbionts can be visualized within reproductive host spore cells

Transmission electron microscopy (TEM) revealed bacterial cells inside developed spores from representative *Amoebophilus,* Chlamydiae, and *Procabacter* host isolates (Fig. 3). Symbiont cells were detectable in the majority of symbiont host spores visualized by TEM. Each symbiont appeared to have gram-negative type cell walls but displayed unique morphologies within spore cells. *Amoebophilus* endosymbionts were rod-like, measured 0.3-0.5µm in width and 0.5-1.3µm in length, and appeared to be distributed throughout the cytoplasm within a host membrane. *Procabacter* endosymbionts were rod shaped, measured 0.25-0.5µm in width and 0.75-1.5µm in length, and in many cases were surrounded by a host membrane. Chlamydiae endosymbionts appeared as dense wrinkled spheres measuring 0.3-0.6µm that were distributed throughout the cytoplasm.

**Figure 3.**
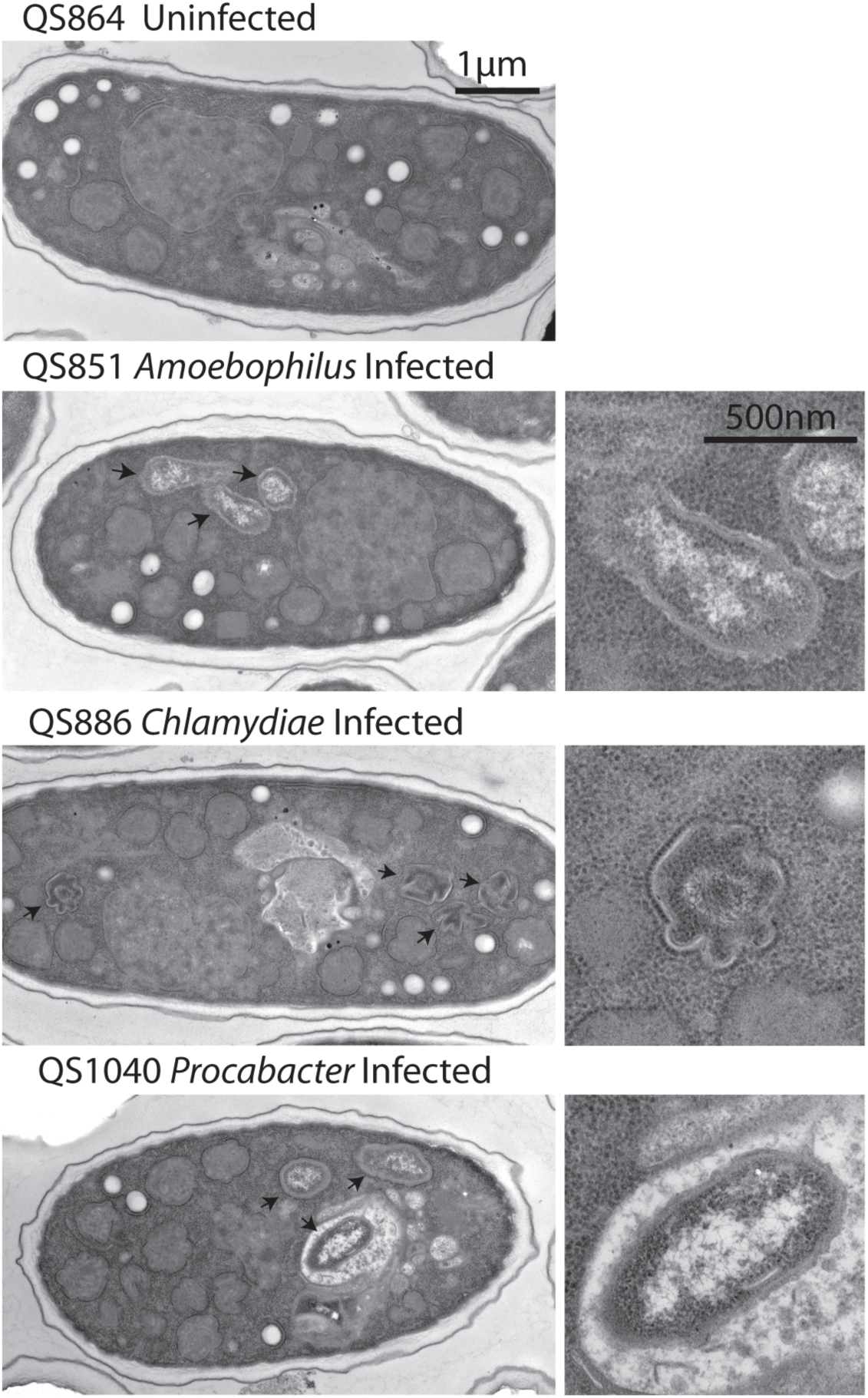
Unculturable symbionts have distinct morphologies in infected spores. Transmission electron micrographs of spores from the indicated representative uninfected and infected *D. discoideum* isolates. Images on the right show close-ups of intracellular bacteria.

### *Amoebophilus* and Chlamydiae can be eliminated from host isolates without significantly impacting host fitness

We did not observe any obvious defects in the growth and development of Chlamydiae and *Amoebophilus* infected hosts. To better assess the impact of these endosymbionts on their hosts, we measured the reproductive fitness of representative host isolates before and after endosymbiont curing (Fig. 4). We eliminated Chlamydiae and *Amoebophilus* from four of their natural hosts through culturing for two social cycles on antibiotic saturated plates. After antibiotic exposure and resumption of development on normal media, we confirmed stable loss of endosymbionts via endosymbiont-specific PCR screening (Fig. 4a). We next cultured 1×10^5^ spores from antibiotic-treated and untreated *D. discoideum* counterparts on nutrient media with *Klebsiella pneumoniae* food bacteria. Following culmination of fruiting body development after a week of incubation, we harvested and measured total spore contents (Fig. 4b). We found no significant differences in spore productivity according to endosymbiont status (one-way analysis of variance, (F(5,42)=1.106, p=0.372). Thus, symbiont infections in these natural host isolates have no significant impact on host spore productivity under these conditions. We could not assess the fitness impact of Chlamydiae or *Amoebophilus* in new host amoebae as our attempts to establish infections in new hosts have been thus far unsuccessful.

**Figure 4.**
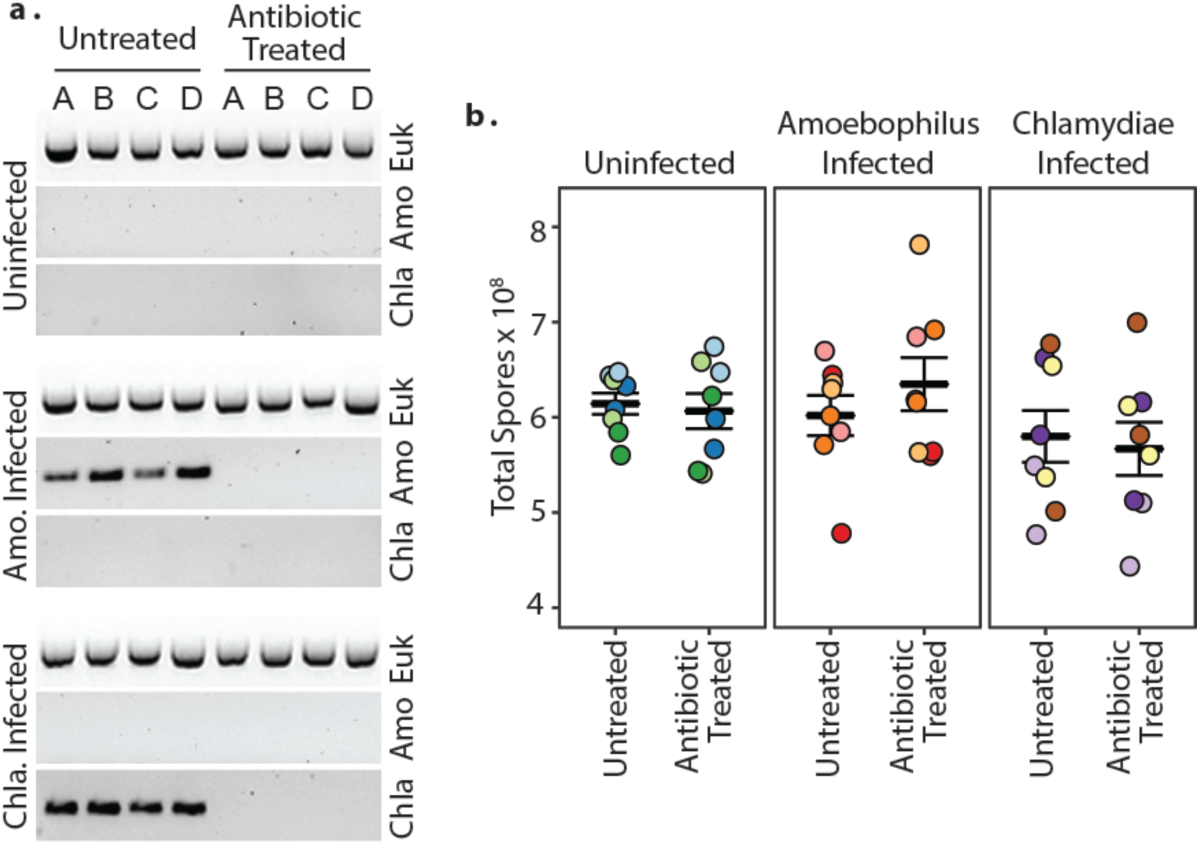
Elimination of *Amoebophilus* and Chlamydiae from natural hosts with antibiotics does not impact host reproductive fitness. Four uninfected, *Amoebophilus*-infected, and Chlamydiae-infected representative host isolates were treated with rifampicin to eliminate symbionts (a) and reproductive fitness (b) was assessed. Symbiont presence is indicated by amplification with *Amoebophilus*-specific or Chlamydiae-specific PCR and DNA gel electrophoresis (with Eukaryote specific gene amplification serving as an internal control) (a). Reproductive fitness was assessed by quantifying total spore productivity of *D. discoideum* cultures after completion of one social cycle (b). Individual *D. discoideum* isolates are represented by point color.

### *Amoebophilus* and Chlamydiae infections do not alter the fitness of hosts during exposure to *Paraburkholderia*

Since co-infections with unculturable symbionts and *Paraburkholderia* occur in the wild, we wanted to determine whether these co-infections would result in a different fitness outcome than *Paraburkholderia* infections alone. We tested the fitness impact of Chlamydiae and *Amoebophilus* during *Paraburkholderia* exposure using the same strategy as above, but with the addition of 5% by volume of the indicated *Paraburkholderia* strain to food bacteria prior to plating (Fig. 5). These *Paraburkholderia* symbionts range from highly detrimental (*P. hayleyella* strain 11), to intermediately detrimental (*P. agricolaris* strains 159 and 70), to neutral (*B. bonniea* strain 859) under the culturing conditions used for this assay (Haselkorn et al., 2018; Khojandi et al., 2019; Shu et al., 2018). We again observed that exposure to distinct *Paraburkholderia* strains significantly influenced spore productivity for both *Amoebophilus* (F(3,60)=36.87, p=<0.001) and Chlamydiae (F(3,60)=28.52, p=<0.001) originating host lines (Fig. 6). However, as with single unculturable symbiont infections, we found that presence or absence of *Amoebophilus* or Chlamydiae had no significant impact on spore productivity during exposure to *P. hayleyella-*11 (F(5,42)=1.171, p=0.339), *B. agricolaris*-159 (F(5,42)=0.985, p=0.438), *P. agricolaris*-70 (F(5,42)=1.095, p=0.377), or *P. bonniea-859* (F(5,42)=0.661, p=0.655) (Fig. 5). Thus, under these conditions the fitness impact of *Paraburkholderia* symbiosis on host spore productivity does not appear to be altered (for better or worse) by co-infections with obligate endosymbionts.

**Figure 5.**
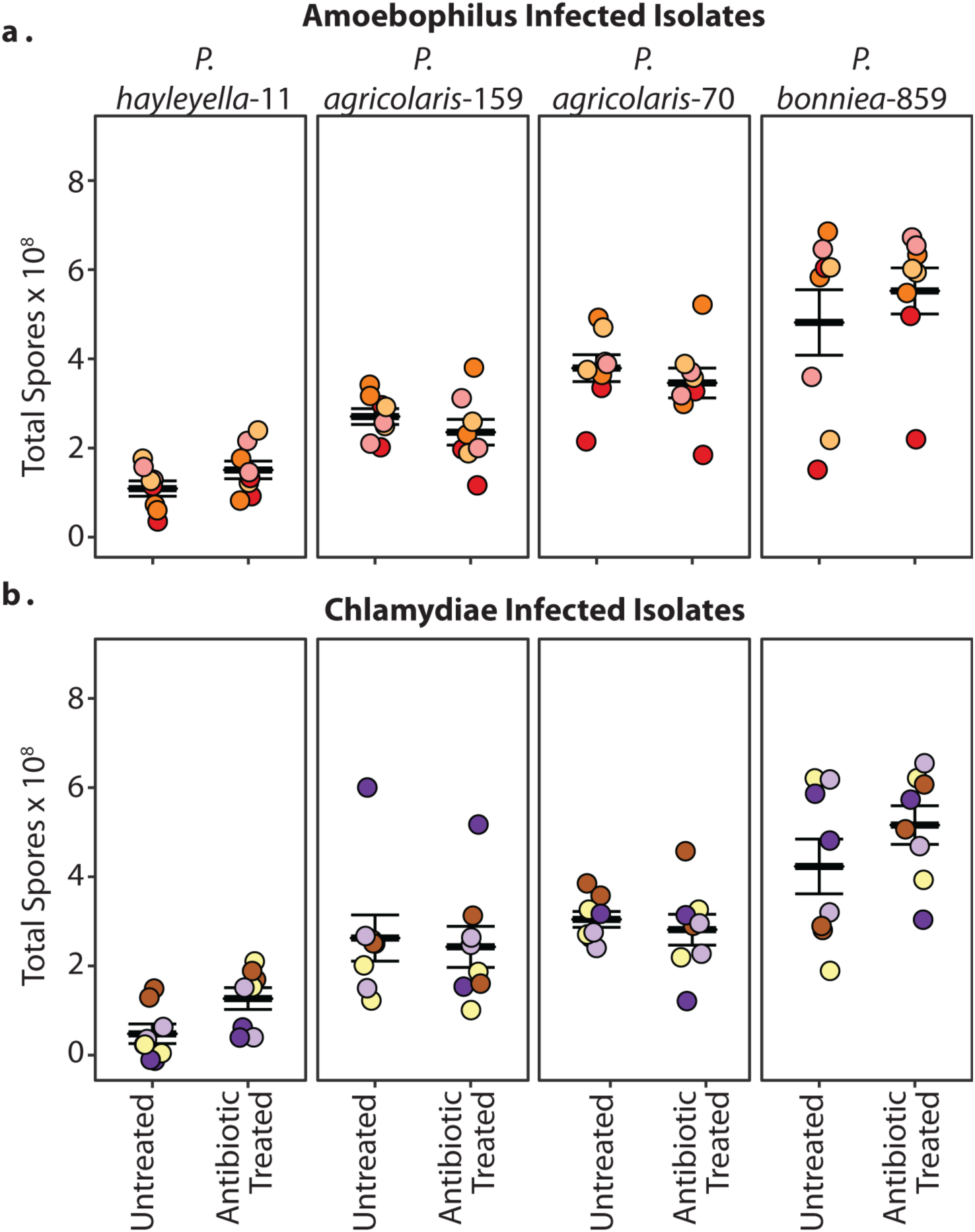
*Amoebophilus* and Chlamydiae infections do not impact the reproductive fitness of hosts co-infected with *Paraburkholderia* symbionts. Reproductive fitness was quantified for four untreated and antibiotic-cured *Amoebophilus*-infected (a) and Chlamydiae-infected isolates (b) following social cycle completion after exposure to the indicated *Paraburkholderia* symbiont strains. Individual *D. discoideum* isolates are represented by point color.

### Detection of positive associations between Chlamydiae and *Paraburkholderia* and *Amoebophilus* and *Paraburkholderia* and a negative association between *Amoebophilus* and Chlamydiae

Our earlier culture-based screening of this *D. discoideum* natural collection found that 25% of isolates were infected with *Paraburkholderia* symbionts (Brock et al., 2020; Haselkorn et al., 2018). For this new round of sequence-based screening, we identified multiple instances of *Paraburkholderia-Amoebophilus* and *Paraburkholderia-*Chlamydiae co-infections in addition to the *Amoebophilus-*Chlamydiae co-infections from above. Specifically, 42 isolates (6%) were co-infected with *Paraburkholderia* and *Amoebophilus*, 25 (3.5%) with *Paraburkholderia* and Chlamydiae, and 9 (1%) were co-infected with all three symbiont types (Fig. 1b). Co-infection prevalence of the different endosymbionts were tested in our three highest sampled populations: Texas-Houston Arboretum, Virginia Mountain Lake 2000, and Virginia Mountain Lake 2014. Pairwise Fisher Exact tests showed a small positive association between Chlamydiae and *Paraburkholderia* in the Texas population (p=0.0132, 15 observed with 10 expected) but not the Virginia populations. There was a positive association between *Paraburkholderia* and *Amoebophilus* in the Virginia 2000 collection (p=0.000277, 43 observed with 30 expected) and a negative association between Chlamydiae and *Amoebophilus* in the Virginia 2014 collection (p=0.000024, none observed with 10 expected).

## Discussion

Many microorganisms, particularly those that are obligately host-associated, are often overlooked because they are unculturable in standard laboratory conditions. The presence of obligate bacterial symbionts in free-living amoebae has been known for some time, particularly for *Acanthamoeba* (Fritsche et al., 1993; Hall and Voelz, 1985; Jeon and Jeon, 1976; Proca-Ciobanu et al., 1975). The list of amoebal endosymbionts and their host species has since expanded but obligate endosymbionts have not been previously described in the social amoeba *D. discoideum* (Corsaro et al., 2010; Delafont et al., 2015; Horn and Wagner, 2004; Samba-Louaka et al., 2019; Schmitz-Esser et al., 2008; Schulz et al., 2014). Here, we screened a collection of 730 *D. discoideum* isolates using a PCR detection strategy in order to identify bacterial associates that could be missed using conventional culturing techniques. Furthermore, to our knowledge, this is the largest collection of amoebae from natural populations to be screened, facilitated by the relative ease in which social amoeba fruiting bodies can be sampled. Here, we find that obligate symbionts are quite common among *D. discoideum* collected from the wild, with bacterial sequences of unculturable bacteria present in 301 isolates (41.2%). All but two of these unculturable bacterial sequences aligned most closely to *Amoebophilus* or *Chlamydiae* species. The identification of these bacteria in *D. discoideum* reveals that this valuable eukaryotic model organism is a natural vector of environmental bacteria. This opens a new role for *D. discoideum* as a research tool for the investigation of how bacterial endosymbionts evolve to live inside hosts, how they affect the fitness of their amoeba host, and how they interact with other bacteria in the amoeba microbiome.

The intracellular morphology of endosymbionts in *D. discoideum* spores were in some cases similar to previous observations of related endosymbionts in other amoebae. *Amoebophilus* cells in *D. discoideum* spores appeared similar to previous observations of *Amoebophilus* symbionts in *Acanthamoeba sp*. (Horn et al., 2001; Schmitz-Esser et al., 2010). The localization of *Procabacter* endosymbionts within a membrane bound vacuole has also been observed for Candidatus *Procabacter* sp. OEW1 in *Acanthamoeba* spp. host cells (Heinz et al., 2007). However, this differs from a previous report on *Procabacter* symbionts in *Acanthamoeba* trophozoites and cysts wherein bacterial cells were not found to be enclosed in vacuoles (Fritsche et al., 2002). The wrinkled morphology of Chlamydiae cells in *D. discoideum* spores is somewhat reminiscent of the spiny structure of *Chlamydia*-like symbionts in *Acanthamoeba, Naegleria,* and *Hartmannella* host species (Casson et al., 2008; Corsaro and Greub, 2006; Horn et al., 2000). However, these morphological observations may be artifacts of (or exaggerated by) sample preparation prior to TEM and may be better resolved using cryo-electron tomography (Huang et al., 2010; Santarella-Mellwig et al., 2013).

*Chlamydiae*-like organisms typically have two or more morphotypes that correspond to distinct developmental stages. Elementary bodies are the infectious form that can be transmitted extracellularly from cell to cell. Once internalized, elementary bodies differentiate into reticulate bodies that typically replicate within inclusion vesicles, although some environmental Chlamydiae can be found directly in the cytosol (Bayramova et al., 2018; Benamar et al., 2017; Bou Khalil et al., 2017; Collingro et al., 2020; Horn, 2008; Nylund et al., 2018). However, in *D. discoideum* host spores, we only visualize one morphology that does not appear to be in an inclusion but is rather distributed throughout the cytosol. It is unclear what morphotype this represents and whether other morphotypes exist in *D. discoideum* prior to spore formation. It is possible that additional Chlamydiae developmental stages occur in vegetative amoebae and that only one form is favored within metabolically inert host spores.

Out of the 251 amoeba isolates from the 2014 Virginia Mountain Lake collection, we identified only two infected with a bacterium with sequence homology to *Procabacter*, suggesting that *D. discoideum* is not a preferred natural host for this endosymbiont. Yet, this *D. discoideum Procabacter* lineage possibly represents an important link for understanding the evolution and adaptation of *Procabacter* to amoeba hosts, as the phylogenetic placement of the novel *D. discoideum Procabacter* lineage is sister to the rest of the *Procabacter* species infecting *Acanthamoeba.* Comparative genome sequencing of this clade, as well as future research into the impact of these infections on their hosts and more extensive wild isolate screenings will better inform our understanding of the significance of these infections in *D. discoideum*.

We identified multiple *D. discoideum* infected with an *Amoebophilus* species that most closely matched Ca. *Amoebophilus asiaticus*, an endosymbiont originally discovered in *Acanthamoeba* (Horn et al., 2001). Similar to the *D. discoideum Procabacter* symbionts, the *D. discoideum Amoebophilus* symbiont lineage is sister to the rest of the *Amoebophilus* clade. The fact that it groups most closely with a putative *Amoebophilus* symbiont discovered in the genome assembly of *Coremiostelium polycephalum,* formerly called *Dictyostelium polycephalum* (Sheikh et al., 2018) suggests that there may be some host specificity of *Amoebophilus* among social amoebae, although sampling of additional amoeba species would be necessary to elucidate this pattern.

The next most closely related bacteria to the *Amoebophilus* genus are the insect facultative endosymbionts of the *Cardinium* genus (Santos-Garcia et al., 2014). These bacteria infect up to 7% of all insect species and have diverse effects on their hosts ranging from reproductive parasitism to nutritional supplementation and defense against parasites. Comparative genome sequencing has uncovered potential routes of adaptation from free-living marine bacteria to either an insect or amoeba host (Santos-Garcia et al., 2014). Including this additional *Amoebophilus* species in comparative genome analysis could further elucidate this process and highlight potential amoeba species-specific adaptations. *Acanthamoeba* and *Dictyostelium* are evolutionarily distant hosts with different lifestyles, thus specific bacterial adaptations may have evolved.

The global significance of *Amoebophilus* infections is under active investigation. *Amoebophilus* have been associated with coral microbiomes and bovine digital dermatitis lesions, which may simply be an indicator that their host amoebae are present in these samples (Apprill et al., 2016; Choi et al., 2009; Horn et al., 2001; Huggett and Apprill, 2019; Zinicola et al., 2015). In amoebae, the full extent of this bacterium’s affect is unknown. The moderately reduced genome of Ca. *Amoebophilus asiaticus* lacks important metabolic pathways but contains a high number of proteins with eukaryotic domains that are thought to be essential for host cell interactions (Schmitz-Esser et al., 2010). Genes associated with host cell exploitation, such as toxins, antimicrobial factors, and a unique secretion system, suggest that Ca. *Amoebophilus asiaticus* is more parasitic than mutualistic (Böck et al., 2017; Penz et al., 2010; Schmitz-Esser et al., 2010). We did not observe an obvious fitness cost of hosting *Amoebophilus* via our spore productivity assays. However, this measures just one aspect of host fitness in defined laboratory conditions and it remains possible that these symbionts may impact host fitness in important ways at different life stages or under different environmental conditions.

Chlamydiae endosymbionts were the most prevalent in the screened collection, with approximately 27% of *D. discoideum* isolates infected. Environmental Chlamydiae appear to be diverse and ubiquitous and many Chlamydiae endosymbionts associate with a wide range of protozoa and other organisms (Corsaro et al., 2013; Corsaro and Venditti, 2006; Coulon et al., 2012). The incredible genetic diversity within the phylum Chlamydiae suggests ancient interactions with eukaryotes, with estimates that the association is 700 million years old (Greub and Raoult, 2003). The diversity of Chlamydiae, as estimated from various 16S rRNA genome databases and sequence read archives, ranges anywhere from 1157-1483 different families (Collingro et al., 2020). This type of screening, however, does not identify the hosts, so insight into adaptation to particular hosts is limited. In this study we have identified eight novel lineages, potentially representing two novel families, both with closely related haplotypes suggesting ongoing evolution within the *D. discoideum* host. The separation of these novel *D. discoideum* Chlamydiae lineages from the other amoeba lineages suggests a long-standing association and possible co-adaptation. As our attempts to transfer *Amoebophilus* or Chlamydiae endosymbionts to new *D. discoideum* hosts were unsuccessful, we could not probe the potential role of host-symbiont adaptation in fitness outcomes. This strengthens the speculation that these symbionts have been associated with their specific host lines for a long period of time but also raises the question of how they were originally acquired. Other lab experiments have shown that some of these Chlamydiae strains are easily transferred across different host genera while others cannot be used to successfully re-infect even their original host amoebae (Corsaro et al., 2013; Coulon et al., 2012; Okude et al., 2012). A previous transfer of a *Protochlamydia* strain to new hosts resulted in a decreased growth rate, suggesting mutual co-evolution between the endosymbiont and its original host (Okude et al., 2012).

Similar to our results for *Amoebophilus*-infected hosts, we did not find any obvious fitness costs from Chlamydiae infection of *D. discoideum* under standard lab conditions for a subset of *D. discoideum* isolates. This does not, however, rule out the possibility that these symbionts may impart significant impacts on their hosts which may be life-stage, environmental, or genotypically context-dependent (Taylor-Brown et al., 2015). Studies of the impact of *Chlamydiae* endosymbionts on their amoeba hosts suggest that the range of outcomes spans the parasite to mutualist continuum (Hayashi et al., 2012). The high prevalence and widespread distribution of particular haplotypes, like haplotype 1, may suggest ecological relevance. It is unclear whether the abundance of particular endosymbiont haplotypes represents an epidemic of a new genotype or selective sweeps due to beneficial effects on their hosts under particular environmental conditions, as seen in insect symbionts (Cockburn et al., 2013; Himler et al., 2011). Depending on the temperature, one *Parachlamydia* endosymbiont of *Acanthamoeba* can either be lytic or endosymbiotic (Greub et al., 2003). The Chlamydiae endosymbiont S13 does not significantly harm its *Acanthamoeba* host and instead has been shown to protect it from *Legionella pneumophila* infection (Ishida et al., 2014; König et al., 2019; Maita et al., 2018; Matsuo et al., 2009; Okude et al., 2012). A *Protochlamydia* endosymbiont has been shown to increase the migration rate, actin density, and growth rate of its native host but had an adverse effect when transferred to non-native hosts (Okude et al., 2012). Overall, future investigations into the impact of these symbionts on hosts using diverse fitness assays and a range of culture conditions will help to elucidate their significance in infected host amoebae.

At least 11.4% (83 out of 730) of these *D. discoideum* are co-infected with some combination of *Chlamydiae*, *Amoebophilus,* or *Paraburkholderia*. The lack of detectable fitness costs from co-infection with unculturable endosymbionts and *Paraburkholderia* in the laboratory is consistent with random associations found in several populations. In different environments, however, other microbe-microbe interactions may have an effect that favors certain combinations of symbionts, as is seen in aphids (Rock et al., 2018). Chlamydiae occur in co-infections less often than would be expected with *Amoebophilus* in one *D. discoideum* population, consistent with the observation that Chlamydiae interact negatively with the *Legionella* pathogen in *Acanthamoeba*. On the other hand, *Paraburkholderia* and *Amoebophilus* are positively associated in one population. *Paraburkholderia* symbionts of *D. discoideum* have been shown to allow other food bacteria to enter into the amoeba spores (Brock et al 2011, DiSalvo et al 2015), and perhaps a similar positive interaction occurs in this case. This effect may outweigh some of the negative interactions of Chlamydiae with other bacteria, as in one population there was positive association with Chlamydiae and *Paraburkholderia*. *Paraburkholderia* symbionts are easily transferred to new hosts via co-culturing and can be found both intracellularly and extracellularly in amoeba cells and structures (DiSalvo et al., 2015; Haselkorn et al., 2018; Shu et al., 2018). *Paraburkholderia* infections in new hosts may be detrimental or beneficial depending on symbiont genotype and environmental context (Haselkorn et al., 2018; Khojandi et al., 2019). Endosymbiont co-infection is not unique to *D. discoideum,* as previous and alternative isolation methods are allowing for identification of co-infections in *Acanthamoeba* (Heinz et al., 2007; Lagkouvardos et al., 2014).

The growing list of bacterial symbionts of free-living amoebae attests to their attractiveness as an intracellular niche for bacterial symbionts. The similarities between amoebae and animal macrophages, as well as the shared intracellular entrance and survival mechanisms employed by endosymbionts and pathogens, further highlight the potential medical and research relevancy of amoebae-symbiosis systems. The discovery of a high prevalence of infection and novel lineages of common amoebae symbiont clades in the model organism *D. discoideum* opens the door to future research on the ecological significance and mechanisms of amoeba-symbiont interactions.

## Methods

### Bacterial Strains and Cultural Conditions

We used *Klebsiella pneumoniae* throughout as our standard *D. discoideum* food supply. For *Paraburkholderia*-unculturable co-infection experiments, we used the *Paraburkholderia* symbionts *P. hayleyella*-11, *P. agricolaris*-70, *P. agricolaris*-159, and *B. bonniea*-859. We grew bacteria on SM/5 agar plates [2 g glucose, 2 g BactoPeptone, 2 g yeast extract, 0.2 g MgCl_2_, 1.9 g KHPO_4_, 1 g K2HPO_5_, and 15 g agar per liter, premixed from Formedium] at room temperature before resuspending them in KK2 buffer [2.25 g KH2PO_4_ (Sigma-Aldrich) and 0.67 g K2HPO_4_ (Fisher Scientific) per liter]. We then adjusted bacterial suspension to a final OD_600nm_ of 1.5. We plated amoebae spores with 200µl of bacterial suspensions for all culturing conditions.

### Symbiont Screening

Total DNA from *D. discoideum* was harvested by suspending 15-25 sori in a 5% chelex solution with 10µg proteinase K, incubating samples for 240 minutes at 56°C, 30 minutes at 98°C, and removing DNA supernatant from chelex beads. PCR with eukaryotic specific primers was run as a DNA extraction control, and symbiont specific primers were used to detect symbionts in each sample (Table S2) (Corsaro et al., 2002; Ossewaarde and Meijer, 1999). For all primer sets we used a touchdown PCR protocol consisting of 94°C for 30 s, 68°C-55°C for 45 x (dropping 1 degree per cycle) for 14 cycles, and 72°C for 1 min, followed by 94°C, 55°C, 72°C for 20 cycles. To confirm infection we ran PCR samples on a one percent agarose gel with TAE buffer and imaged on a transilluminator. PCR products were sent for sequencing (Genewiz), and the resulting Sanger sequences were cleaned using the program Geneious v8 (http://www.geneious.com (Kearse et al., 2012)). We use NCBI Blast to initially identify the bacterial species/strain based on highest sequence similarity. Sequences have been deposited to GenBank under accession numbers (MT596490-MT596496).

### Amoebae Clone Types and Culture Conditions

Amoebae were cultured from freezer stocks by resuspending frozen spores onto SM/5 agar plates with 200µl of food *K. pneumoniae* suspensions. Plates were incubated at room temperature under low lights and ∼60% humidity for approximately 1 week (or until fruiting body development).

For antibiotic curing and spore quantification, we used four uninfected, four *Amoebophilus* infected, and four Chlamydiae infected host lines. Uninfected representatives were QS864, QS970, QS1003 and QS1069 (highlighted in yellow in Table S1). *Amoebophilus* infected representatives included QS851, QS1002, QS1011, and QS1017 (highlighted in green in Table S1). Chlamydiae infected representatives included QS866, QS991, QS1023, and QS1089 (highlighted in blue in Table S1). For all experiments, we initiated experimental conditions from pre-developed fruiting bodies by suspending ∼8-15 sorus contents in KK2. We then plated 10^5^ spores onto an SM/5 plate with 200µl of bacterial suspensions (*K. pneumoniae* only or with 95% *K. pneumoniae/* 5% *Paraburkholderia* mixtures by volume for co-infection experiments). Plates were then incubated as described above for the time indicated for each experimental condition.

### Curing of Symbiont Infections

We treated uninfected, Chlamydiae-infected, and *Amoebophilus* infected representative isolates with rifampacin by plating ∼10^2^ spores with 400µl *K. pneumoniae* on SM/5 medium containing 30µg/ml rifampacin. Plates were incubated until fruiting body formation. Sori from grown fruiting bodies were then harvested and replated for a second round of antibiotic exposure (using the same strategy as described above). Following fruiting body formation, spores were harvested and transferred to SM/5 with *K. pneumoniae* and incubated. Following fruiting body formation on non-antibiotic plates, we extracted total sorus content DNA and checked for symbiont infection using the Chlamydiae-specific and *Amoebophilus*-specific PCR protocol described above.

### Total Spore Productivity Assay

To determine total spore productivity of antibiotic treated and untreated *D. discoideum* isolates we plated 10^5^ spores on SM/5 agar with 200µl of *K. pneumoniae* and incubated at room temperature. Five days after plating, total plate contents were collected into ∼10ml KK_2_ + NP-40 alternative detergent solution. Solutions were then diluted appropriately (∼20x) and spores quantified via hemocytometer counts. We measured and averaged over at least three replicates. To determine spore productivity after exposure to *Paraburkholderia*, we conducted the assay as described but plated spores with 200µl of a 95/5% by volume mixture of *K. pneumoniae/Paraburkholderia* cell suspensions.

### Statistics

Significant positive or negative associations between co-infecting endosymbionts were determined using a Fisher Exact test (2×2 contingency table) on pairwise comparisons in our three most highly sampled populations. Statistical analyses for spore productivity experiments were performed in R 3.6.0. Comparisons were done using one-way ANOVA’s with antibiotic treatment as the independent variable.

### Transmission Electron Microscopy

Complete processing and imaging of spores by transmission electron microscopy was performed by Wandy Beatty at the Washington University Molecular Microbiology Imaging Facility. *D. discoideum* isolates imaged included QS864, QS851, QS886, QS1040. Briefly, spores were harvested in KK2 buffer and fixed with 2% paraformaldehyde with 2.5% glutaraldehyde (Polysciences Inc., Warrington, PA) in 100 mM cacodylate buffer (pH 7.2) for 1-3 hrs. at room temperature. We washed samples with cacodylate buffer and postfixed in 1% osmium tetroxide (Polysciences Inc.) for 1 hr before rinsing in dH_2_O prior to *en bloc* staining with 1% aqueous uranyl acetate (Ted Pella Inc., Redding, CA) for 1 hr. We dehydrated samples in increasing concentrations of ethanol and embedded them in Eponate 12 resin (Ted Pella Inc.). Sections of 95 nm were cut with a Leica Ultracut UCT ultramicrotome (Leica Microsystems Inc., Bannockburn, IL), stained with uranyl acetate and lead citrate, and viewed on a JEOL 1200 EX transmission electron microscope (JEOL USA Inc., Peabody, MA) equipped with an AMT 8 megapixel digital camera (Advanced Microscopy Techniques, Woburn, MA).

### Phylogenetic Tree Construction

The *Procabacter* full length 16S rRNA phylogeny was created using the top ten NCBI Blast results for the *Procabacter* endosymbiont of D. *discoideum* sequence and representative species from the phylum *Betaproteobacteria*. The species were chosen using the NCBI taxonomy database and 16S rRNA sequences were obtained from the NCBI nucleotide database as well as the three top hits to the SILVA database. A sequence from *Rickettsia rickettsii* was also obtained from the NCBI nucleotide database to serve as an outgroup. Sequences were imported into Mega7 (Tamura et al, 2013) and aligned with MUSCLE. All sequences were then trimmed to a uniform 1396 bp. The Kimura 2-parameter model with a discrete Gamma distribution was selected using the “Find Best DNA/Protein Models” command in Mega7. A Maximum Likelihood tree was then constructed with 1000 bootstrap replicates.

The *Amoebophilus* full length 16S rRNA phylogeny was created using the top five NCBI Blast results for the *Amoebophilus* endosymbiont of *D. discoideum* sequence, a sequence from the recently sequenced *Dictyostelium polycephalum* genome assembly, a sequence from the related endosymbiont *Candidatus Cardinium hertigii*, and representative species from the order *Cytophagales*. The species were chosen using the phylogeny in Hahnke et al., (2016) as a guide, and full length 16S rRNA sequences were obtained from the NCBI nucleotide database (Hahnke et al., 2016) as well as the three top hits from the SILVA database. A sequence from *Bacteroides fragilis* was also obtained from the NCBI nucleotide database to serve as an outgroup. Sequences were imported into Mega7, aligned with MUSCLE, and trimmed to a uniform 1364 bp. The Kimura 2-parameter model with a discrete Gamma distribution was selected using the “Find Best DNA/Protein Models” command in Mega7. A Maximum Likelihood tree was then constructed with 1000 bootstrap replicates.

The Chlamydiae full length 16S rRNA phylogeny was created using 1-2 representative species from the nine families in the phylum Chlamydiae from Taylor-Brown et al., (2015), 1 from *Ca.* Arenachlamydiaceae (Pillonel et al. 2018), and the three top hits for each haplotype from the SILVA database. Full length 16S rRNA sequences of chosen species were obtained from both the NCBI nucleotide and SILVA databases. Sequences were imported into Mega7 and aligned with MUSCLE, and then trimmed to 1354 bp. The Tamura 3-parameter model with a discrete Gamma distribution was selected for the short tree, and the General Time Reversal model with a discrete Gamma distribution and invariant sites was selected for the full-length tree using the “Find Best DNA/Protein Models” command in Mega7. A Maximum Likelihood tree was then constructed for each with 1000 bootstrap replicates.

### Verification of symbiont strain novelty

In order to verify that, to the best of our knowledge, these *D. discoideum* Chlamydiae lineages are novel, we searched the SILVA database (in addition to the NCBI Blast database) for nearest neighbors. A table of the sequence identities to the SILVA database for all of our full length 16S rRNA endosymbiont haplotypes is included as Supplemental Table 2. Two recent papers (Pillonel et al., 2018 and Dharamshi et al., 2020) have newly discovered Chlamydiae lineages so we compared our 16S rRNA sequences to theirs where possible. The phylogenetic analyses from the Pillonel et al. paper were done from metagenomic sequences, which often lack identified 16S rRNA genes, and thus it was not always possible to compare directly. We did use BLAST to query our sequences against these metagenomes in NCBI and in no case did we find a match with >85% sequence identity, if a match was found at all. For the Dharamshi et al. paper we separately aligned our sequences (full length Chlamydiae haplotypes 1-3) with their publicly available 16S rRNA sequences. We ran a Neighbor-joining phylogenetic analysis in MEGA 7 and computed a pairwise distance matrix. Finally, we searched the IMNGS database (Lagkouvardos et al. 2016), which is designed to query sequence read archives from NCBI for 16S rRNA sequences, for matches to our most common *D. discoideum* Chlamydiae haplotype.

## Acknowledgements

We thank members of the Haselkorn, DiSalvo, and Strassmann-Queller labs for their support and suggestions, in particular Debbie Brock. We also thank the many people in the group that collected the samples, in particular Debbie Brock, Tom Platt and Angelo Fortunato. We also thank Dierdra Renfroe for her attempts to transfer *Chlamydiae* symbionts to new host amoebae. This material is based upon work supported by the National Science Foundation under grant numbers DEB-1753743 and IOS-1656756 to JES and DCQ.

---

**Table S1. Table of Screened D. discoideum Natural Isolates.** Table lists each *D. discoideum* isolate by identification number and location of collection. PCR amplification of *Dictyostelium*, *Paraburkholderia*, *Amoebophilus*, or Chlamydiae specific DNA is indicated with a 1 (for positive amplification) or 0 (for negative amplification). Samples used for fitness assays are highlighted in yellow.

**Table S2. SILVA Database Classification of Symbiont Haplotypes.** Percent sequence identity to closest hit in the database for the five full length 16S rRNA sequence haplotypes and their least common ancestor taxonomic classification.

**Table S3. IMNGS database >99% matches for Chlamydiae Endosymbiont of *D. discoideum* Haplotype 1.** Includes metagenome sample type and collection location.

**Table S4. PCR Primers Used for this Study.** PCR primer names, sequences, and associated references.

