## Supplementary figures and images for "Novel Chlamydiae and *Amoebophilus* endosymbionts are prevalent in wild isolates of the model social amoeba *Dictyostelium discoideum*"

### Supplemental Table 4

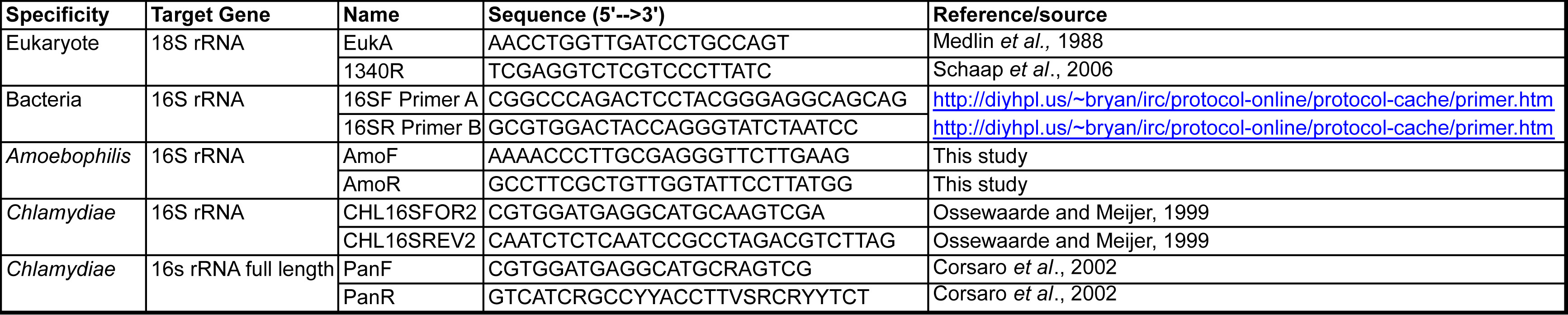
